# Cross-session generalization in automated behavioral tracking of *Galleria mellonella* larvae: comparison of classical computer vision, deep learning and generative domain adaptation

**DOI:** 10.64898/2026.07.24.740557

**Authors:** Kasjan Śmigielski, Natalia Piórkowska

## Abstract

**Background:** Automated behavioral tracking is increasingly used in biological and biomedical research; however, robustness across heterogeneous imaging conditions remains a major challenge. Domain shifts caused by changes in illumination, contrast, or acquisition setup can substantially degrade the performance of computer vision and deep learning models, limiting their practical applicability. This issue is particularly relevant for small-scale biological datasets, where extensive annotation and retraining are often impractical.

**Methods:** We developed a behavioral tracking framework for *Galleria mellonella* larvae combining a classical computer vision (CV) pipeline, a YOLOv8s-seg + ByteTrack deep learning pipeline, and generative domain adaptation methods. Behavioral recordings were collected in two independent experimental sessions under distinct illumination conditions (top and bottom lighting), creating a natural cross-session domain shift. The classical pipeline was based on contour detection, adaptive preprocessing, temporal smoothing, and trajectory reconstruction. Deep learning models were trained on 320 manually annotated frames and evaluated in both within-session and cross-session settings. To improve generalization, we investigated generative augmentation using StyleGAN2-ADA and unpaired image-to-image translation using CycleGAN. Tracking outputs were further analyzed through behavioral descriptors, including trajectories, traveled distance, velocity, spatial occupancy heatmaps, and directional movement patterns.

**Results:** The classical CV pipeline achieved high detection performance in both recording sessions, with mean detection rates of 99.05% and 99.71%, respectively. A YOLOv8s-seg model demonstrated strong within-session performance but exhibited severe degradation under cross-session evaluation, with larval mask segmentation performance dropping to mAP@0.5 = 9.1%, confirming the presence of a substantial domain shift. Despite differences in detection methodology, behavioral metrics derived from YOLO and CV pipelines showed strong agreement at the group level (Pearson correlation r = 0.89; median distance ratio = 0.99). Generative augmentation with StyleGAN2-ADA did not yield meaningful gains in tracking robustness — likely because baseline performance was already near ceiling — whereas CycleGAN-based domain adaptation substantially reduced the domain gap and improved cross-session detection performance while preserving biologically relevant trajectory structures and spatial behavioral patterns.

**Conclusions:** Cross-session variability represents a critical challenge for automated behavioral tracking in biological experiments. Our results demonstrate that carefully designed classical computer vision approaches can achieve highly reliable tracking in small-data settings, while deep learning models require explicit strategies to address domain shift. Generative domain adaptation, particularly CycleGAN-based image translation, offers an effective solution for improving cross-session generalization without additional manual annotation. The proposed framework provides a robust foundation for scalable behavioral phenotyping of *Galleria mellonella* and other small biological model organisms.

## 1. Introduction

Automated behavioral analysis has become an indispensable component of contemporary biological and biomedical research, enabling objective, high-throughput, and reproducible quantification of animal phenotypes. Recent advances in computer vision (CV) and deep learning (DL) have transformed the study of animal behavior by providing automated tools for tracking movement and extracting spatiotemporal features across a wide range of model organisms (Mathis et al., 2018; Pereira et al., 2022). The resulting descriptors — trajectories, locomotor activity, spatial occupancy, and exploration strategies — are increasingly used as phenotypic biomarkers in physiology, toxicology, pharmacology, and infectious disease research.

Despite this progress, the robustness and generalizability of behavioral tracking systems remain major challenges. In practice, image acquisition conditions often vary across recording sessions due to differences in illumination, camera positioning, background, or hardware. Such variations induce domain shifts between datasets, reducing segmentation accuracy and introducing systematic errors in downstream behavioral measurements. Pipelines that perform excellently under controlled conditions frequently deteriorate on data acquired under different imaging settings, making cross-session generalization a critical requirement for real-world deployment.

This problem is especially acute in small-data biological applications, where large annotated datasets are impractical. Deep learning detectors such as YOLO (Redmon et al., 2016; Jocher et al., 2023) rely on extensive annotated data and may generalize poorly to distributions absent from training (Wang et al., 2025); transfer learning and augmentation only partially alleviate this. Classical computer vision, by contrast, requires fewer data and offers greater interpretability, but depends on session-specific parameter tuning that limits scalability and reproducibility.

The larvae of *Galleria mellonella* are an increasingly important invertebrate model in biomedical research (Tsai et al., 2016; Pereira et al., 2018). Their low maintenance cost, ease of handling, and physiological relevance to infection and toxicity studies make them a popular alternative to vertebrate models (Wojda, 2017; Loh et al., 2013), and their behavioral changes have been linked to stress, infection, and bioactive compounds, rendering locomotor activity an informative phenotypic readout (Adamo, 2006; Dantzer, 2001). Yet, compared with *Drosophila melanogaster*, zebrafish, or rodents, computational frameworks for automated behavioral analysis of *G. mellonella* remain underdeveloped, lacking standardized tracking pipelines and systematic robustness evaluations across heterogeneous imaging conditions.

In this study, we develop and evaluate a behavioral tracking framework for *Galleria mellonella* larvae that combines classical computer vision, deep learning-based detection, and generative domain adaptation. Two independent recording sessions acquired under substantially different illumination conditions serve as a natural domain-shift scenario. We (i) design and validate a robust classical CV pipeline for larval tracking and feature extraction; (ii) implement a YOLOv8s-seg + ByteTrack workflow and evaluate it under within- and cross-session conditions; (iii) investigate two complementary generative strategies — StyleGAN2-ADA augmentation to enrich the limited training set and unpaired CycleGAN domain adaptation to mitigate cross-session degradation; and (iv) assess how tracking accuracy propagates to downstream behavioral metrics, including trajectories, traveled distance, velocity, spatial occupancy, and heatmap-based representations.

The main contributions of this study are as follows:

1. Development of a robust and interpretable computer vision framework for automated tracking of *Galleria mellonella* larvae.
2. Systematic quantification of cross-session domain shift in biological behavioral tracking under heterogeneous illumination conditions.
3. Comparative evaluation of classical computer vision and deep learning approaches in a small-data experimental setting.
4. Investigation of StyleGAN2-ADA-based generative augmentation and CycleGAN-based domain adaptation as strategies for improving tracking robustness.
5. Demonstration of the impact of tracking performance on biologically meaningful downstream behavioral measurements.

By integrating classical computer vision, deep learning, and generative domain adaptation within a unified framework, this work provides new insights into the challenges of behavioral tracking under realistic acquisition variability and offers practical strategies for improving robustness and reproducibility in automated phenotyping studies.

## 2. Related Work

### 2.1. Automated Tracking of Animal Behavior

Automated behavioral tracking is a cornerstone of quantitative biology. Early systems relied on classical computer vision — background subtraction, thresholding, contour extraction (Suzuki and Abe, 1985; Canny, 1986), centroid estimation, and Kalman filtering — often delivered through commercial platforms such as EthoVision (Noldus et al., 2001). These approaches remain widely used for their efficiency, interpretability, and minimal training requirements (Bradski and Kaehler, 2008), especially in controlled environments with stable appearance and background.

Deep learning (LeCun et al., 2015; Goodfellow et al., 2016) has greatly expanded these capabilities, enabling robust detection, instance segmentation, pose estimation, and multi-object tracking under challenging conditions. Systems such as DeepLabCut (Mathis et al., 2018), SLEAP (Pereira et al., 2022), idtracker.ai, and YOLO-based pipelines (Redmon et al., 2016; Jocher et al., 2023) achieve high accuracy across rodents, zebrafish, insects, and non-model organisms (Wang et al., 2025; Manduca et al., 2025), yielding detailed descriptors of locomotion, posture, and movement dynamics. However, these models typically require substantial annotated data and are sensitive to distribution shifts between training and deployment, so robust generalization remains a major challenge in biological applications with limited annotation resources and variable acquisition conditions.

### 2.2. Behavioral Tracking in Invertebrate and Small-Scale Biological Models

Automated behavioral analysis has been extensively developed for established organisms such as *Drosophila melanogaster* (Sokolowski, 2001), *Caenorhabditis elegans* (Wang et al., 2025), and zebrafish larvae, supported by dedicated tools that quantify locomotion, exploration, and responses to perturbations. For zebrafish larvae, deep learning pose-estimation and tracking pipelines have been built on DeepLabCut- and SLEAP-derived frameworks (Lau et al., 2023; Li et al., 2025), and synthetic training data have enabled markerless three-dimensional pose estimation (Johnson et al., 2023). For *Drosophila* larvae, tools such as the Hatching-Box (Pegoraro et al., 2024) and LarvaTagger (Carreira-Rosario et al., 2024) illustrate the breadth of available larval phenotyping.

By comparison, few studies address automated tracking of invertebrate larvae used in biomedical and infection research (Brackenbury, 1997; Manduca et al., 2025). Small organisms pose specific challenges — limited visual features, low resolution, variable morphology, and sensitivity to imaging artifacts — under which carefully designed classical CV often remains competitive with deep learning, particularly in controlled environments where organisms can be spatially isolated.

The greater wax moth larva, *Galleria mellonella*, has become an important model in microbiology, immunology, toxicology, and antimicrobial drug discovery (Tsai et al., 2016; Pereira et al., 2018; Wojda, 2017; Loh et al., 2013), with behavioral responses linked to infection, stress, and bioactive compounds (Adamo, 2006; Dantzer, 2001). Nevertheless, computational methodologies for this organism remain underdeveloped: existing studies emphasize biological outcomes rather than tracking accuracy, robustness, and reproducibility, and standardized frameworks or benchmarks for *G. mellonella* are largely absent.

### 2.3. Domain Shift in Biological Imaging and Behavioral Analysis

A critical yet frequently overlooked challenge is domain shift — a discrepancy between the statistical distributions of training and testing data. In biological imaging it arises from variations in illumination, camera configuration, optical setup, resolution, background, or protocol, and even minor changes in acquisition conditions can substantially degrade segmentation, detection, and trajectory reconstruction. This is particularly relevant in longitudinal and multi-session experiments collected over time with slightly different settings.

Although domain shift is well studied in medical imaging and general computer vision, systematic evaluations of cross-session robustness remain uncommon in behavioral phenotyping, and few studies examine how domain-shift-induced tracking errors propagate to downstream measurements such as traveled distance, velocity, spatial occupancy, or exploratory indices. Improving cross-session generalization therefore remains an important open methodological challenge.

### 2.4. Generative Models for Data Augmentation and Domain Adaptation

Generative artificial intelligence offers powerful strategies for limited data and domain mismatch. Generative adversarial networks (GANs; Goodfellow et al., 2014) excel at image synthesis, style transfer, and augmentation. Synthetic image generation is increasingly used to expand training datasets when annotation is costly, and GAN-generated images have improved segmentation, classification, and detection in biomedical imaging; StyleGAN2-ADA in particular performs well in small-data regimes via adaptive discriminator augmentation, which stabilizes training and reduces overfitting (Karras et al., 2020).

Beyond synthesis, domain adaptation reduces discrepancies between image distributions. CycleGAN (Zhu et al., 2017) is among the most influential unpaired image-to-image translation methods, transforming images between acquisition domains while preserving structural content — applied successfully in medical imaging, microscopy, histopathology, and industrial inspection. By harmonizing appearance across conditions, it reduces domain gaps without additional annotation. Despite these advances, generative augmentation and domain adaptation remain little explored for behavioral tracking, and it is largely unknown to what extent they improve cross-session generalization while preserving biologically meaningful trajectory information.

### 2.5. Research Gap and Study Rationale

Despite progress in both classical and deep learning-based tracking, several gaps remain. First, the impact of cross-session domain shift on tracking accuracy and downstream behavioral measurements has rarely been quantified systematically, despite being a major obstacle to routine deployment. Second, dedicated computational frameworks for *Galleria mellonella* are largely lacking, even though reliable locomotor phenotyping of this widely used infection and toxicity model could serve as a non-invasive biomarker of infection severity and compound toxicity. Third, the effectiveness of generative augmentation and domain adaptation for behavioral tracking under heterogeneous imaging conditions — and whether such methods preserve biologically meaningful information after image transformation — remains insufficiently understood.

The present study addresses these gaps by integrating classical computer vision, deep learning-based detection and tracking, generative augmentation, and CycleGAN-based domain adaptation within a unified framework. Using *Galleria mellonella* larvae recorded under distinct illumination conditions as a natural domain-shift scenario, we systematically evaluate tracking performance, cross-session generalization, and the influence of tracking accuracy on downstream behavioral metrics, aiming to inform the development of robust, reproducible phenotyping pipelines for small biological model organisms.

## 3. Methods

### 3.1. Overview of the Framework

The proposed framework consisted of a multi-stage pipeline for automated behavioral analysis of *Galleria mellonella* larvae. Starting from raw video recordings containing six simultaneously recorded Petri dishes, the workflow included dish isolation, image preprocessing, larval detection, trajectory reconstruction, temporal stabilization, behavioral feature extraction, and quantitative evaluation.

To investigate robustness across heterogeneous acquisition conditions, three methodological approaches were evaluated: (i) a classical computer vision (CV) tracking pipeline, (ii) a deep learning-based detection and tracking pipeline based on YOLOv8s-seg and ByteTrack, and (iii) generative methods aimed at improving cross-session generalization through StyleGAN2-ADA-based augmentation and CycleGAN-based domain adaptation.

The complete workflow was designed to assess not only tracking accuracy but also the impact of detection performance on downstream behavioral metrics derived from larval movement trajectories.

### 3.2. Dish Isolation and Spatial Calibration

Behavioral recordings were acquired in two independent experimental sessions conducted on different dates and under distinct illumination conditions: Session I (top LED illumination) and Session II (bottom LED illumination). Both sessions used identical acquisition hardware (Canon XF300 camera, 1920 × 1080 px, 25 frames per second, H.264/MP4). In each session, larvae were divided into five experimental groups, with six larvae per group (one larva per Petri dish). Each group was recorded in a single video containing all six dishes simultaneously, yielding 5 videos per session and 10 videos in total (60 larva-recordings overall, 30 per session). Recording duration was approximately 30 minutes per group in Session I and 15 minutes per group in Session II, corresponding to roughly 45,000 and 22,500 frames per video, respectively. Recording began after a 5-minute acclimatization period.

Video recordings contained six Petri dishes arranged in a 2 × 3 configuration. To enable independent analysis of individual larvae, each recording was divided into six fixed regions of interest (ROIs) corresponding to individual dishes. Dish locations were manually defined in the initial frame and subsequently applied to all frames within a recording.

Spatial calibration was performed using the known Petri dish diameter of 95 mm. Pixel coordinates were converted into physical units using a constant scaling factor determined during calibration. Based on measurements across recordings, a scale factor of 0.20 mm/pixel was adopted for all subsequent analyses.

Each ROI contained a single larva, ensuring that tracking was performed under a single-object scenario.

### 3.3. Classical Computer Vision Tracking Pipeline

#### 3.3.1. Image Preprocessing

Each ROI was converted to grayscale and processed independently. Gaussian filtering with a 7 × 7 kernel (selected empirically among 3 × 3, 5 × 5, and 7 × 7 configurations) was applied to suppress high-frequency noise and improve segmentation stability. To restrict analysis to the interior of the Petri dish, a dish mask with rounded corners was applied, eliminating reflections and edge artifacts originating from dish boundaries.

#### 3.3.2. Session-Specific Processing

Two recording sessions were acquired under different illumination conditions.

Session I employed top illumination, producing a relatively dark background and bright larval appearance. In this case, grayscale conversion, thresholding, and morphological processing were sufficient for reliable segmentation.

Session II employed bottom illumination, resulting in a bright background and darker larval appearance. To compensate for reduced local contrast and illumination variability, Contrast Limited Adaptive Histogram Equalization (CLAHE) was introduced before thresholding. All classical image-processing operations were implemented using the OpenCV library (Bradski and Kaehler, 2008), with contour extraction based on the border-following algorithm of Suzuki and Abe (1985).

These session-specific preprocessing procedures provided an interpretable example of domain shift between acquisition conditions.

#### 3.3.3. Segmentation and Morphological Processing

Foreground segmentation was performed using intensity thresholding. Threshold values were optimized separately for each session to account for differences in image contrast. To suppress noise and improve contour continuity, morphological erosion and dilation operations were applied sequentially. The order and parameters of these operations were empirically optimized to preserve larval morphology while minimizing false-positive detections.

#### 3.3.4. Contour Selection and Tracking

Contours extracted from binary masks were filtered according to geometric and temporal constraints.

Candidate contours were evaluated using:

- contour area,
- distance from the previously detected position,
- temporal consistency.

To reduce susceptibility to artifacts, an area-penalty scoring mechanism was introduced. This approach balanced contour size and spatial proximity, favoring biologically plausible detections while suppressing reflections and fragmented structures.

When no contour satisfied selection criteria, the largest available contour was retained as a fallback solution.

#### 3.3.5. Temporal Stabilization

Centroid positions were smoothed using an exponential moving average

(EMA) filter with α = 0.3.

Missing detections were reconstructed by linear interpolation between neighboring valid positions. This procedure reduced frame-to-frame fluctuations while preserving biologically relevant movement patterns.

### 3.4. Behavioral Feature Extraction

Behavioral descriptors were computed from centroid trajectories.

Let (*x_t_*, *y_t_*) denote the larval centroid position at frame *t*, *T* the total number of frames, *FPS* = 25 the recording frame rate, and *s* = 0. 20 *mm*/*px* the spatial scaling factor.

Pixel-to-millimeter conversion:

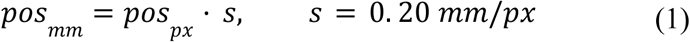

The total traveled distance was calculated as:

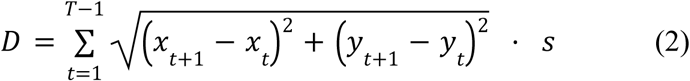

Average velocity was calculated as:

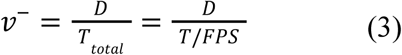

where *D* is the total traveled distance and *T_total_* denotes recording duration. Spatial coverage was defined as the proportion of unique locations visited by the larva within the Petri dish area:

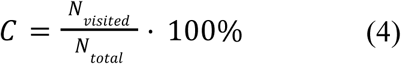

where *N_visited_* is the number of spatial bins visited by the larva and *N_total_* is the total number of bins covering the Petri dish area. Movement direction angle (compass convention):

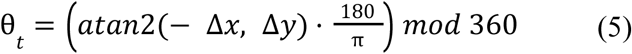

where Δ*x* = *x_t_*_+1_ – *x_t_* and Δ*y* = *y_t_*_+1_ − *y_t_*

Additional behavioral descriptors included:

- movement trajectories,
- spatial occupancy heatmaps,
- spatial coverage.

Together, these features provided a quantitative representation of larval locomotor behavior.

### 3.5. Deep Learning-Based Detection and Tracking

#### 3.5.1. Annotation Dataset

A total of 320 video frames were manually annotated using Roboflow. The dataset included 160 frames from Session I and 160 frames from Session II. Frames were sampled approximately every 100 frames (about every 4 s at 25 fps) from the full recordings, ensuring coverage of different experimental groups, larval positions (center, edge, near the dish wall), and time points throughout each recording.

Two object classes were defined:

1. larva – instance segmentation polygons,
2. dish – bounding-box annotations.

Each frame contained six larvae and six Petri dishes.

The annotated dataset was partitioned into training (70%, 224 frames), validation (20%, 64 frames), and test (10%, 32 frames) subsets. Because frames were drawn from a small number of long videos (5 per session), the split was performed at the frame level rather than at the video level; the implications of this design for potential information leakage are addressed explicitly in Section 3.5.4. In addition, session-specific dataset versions were generated to enable the cross-session experiments described below.

#### 3.5.2. YOLOv8 Instance Segmentation

Instance segmentation was performed using YOLOv8s-seg (Jocher et al., 2023), a single-stage detector building on the YOLO family of real-time object detectors (Redmon et al., 2016).

Training employed transfer learning initialized from pretrained COCO weights. The dataset was divided into training, validation, and test subsets according to the experimental design described below.

Model performance was assessed using standard object detection and segmentation metrics, including precision, recall, and mAP.

#### 3.5.3. Multi-Object Tracking

Predicted larval masks were linked across frames using ByteTrack (Zhang et al., 2022). Tracking was performed on full-frame recordings without prior dish isolation, enabling simultaneous tracking of all larvae present in the recording. Trajectories reconstructed by ByteTrack were subsequently transformed into dish-centered coordinates and analyzed using the same behavioral metrics as the classical CV pipeline.

#### 3.5.4. Data Splitting and Information Leakage

Because annotation frames were extracted from a limited number of long recordings, the train/validation/test partition was necessarily performed at the frame level rather than at the video level. We therefore explicitly considered the risk of information leakage, i.e., the possibility that highly similar frames from the same video could be distributed across the training and evaluation subsets, which could inflate within-session performance estimates. Two design features mitigate, but do not entirely eliminate, this risk for the within-session evaluation. First, annotation frames were temporally subsampled at intervals of approximately 100 frames (≈4 s); given the slow locomotion of the larvae, consecutive sampled frames still differ in larval pose and position, although they share the same static background, dish geometry, and illumination. Consequently, the within-session metrics (Table 2) may be subject to a positive bias arising from the background and acquisition similarity between the training and test frames of the same session, and should be regarded as an upper bound on within-session performance.

Crucially, however, the central finding of this study — the cross-session evaluation (Table 3) — is structurally free of this form of leakage. In the cross-session experiments, the model is trained exclusively on frames from one session and evaluated exclusively on frames from the other, physically independent recording session acquired on a different date and under different illumination. There is no overlap of videos, backgrounds, or acquisition conditions between the training and test domains. The severe cross-session degradation reported below (larval mask mAP@0.5 dropping to 9.1% for S1→S2 and to 0.007% for S2→S1) therefore cannot be attributed to information leakage. On the contrary, any residual positive bias in the within-session estimates would render the observed cross-session gap a conservative estimate of the true domain-shift effect.

### 3.6. Generative Augmentation

#### 3.6.1. StyleGAN2-ADA Training

To investigate whether synthetic data could improve robustness in small-data conditions, StyleGAN2-ADA (Karras et al., 2020) was trained using image crops extracted from annotated recordings.

Adaptive discriminator augmentation was employed to stabilize training and reduce overfitting associated with limited training data.

#### 3.6.2. Synthetic Dataset Generation

After convergence, the trained generator was used to create synthetic Petri dish images containing larval appearances representative of the training domain.

Generated samples underwent visual quality control and automatic pseudo-labeling using a pretrained YOLOv8s-seg model. Low-quality synthetic images were discarded prior to inclusion in downstream experiments.

#### 3.6.3. Evaluation Strategy

Models trained with and without synthetic augmentation were compared to assess the effect of generated data on within-session and cross-session performance.

### 3.7. CycleGAN-Based Domain Adaptation

#### 3.7.1. Definition of Domains

Two image domains were defined:

- Domain A: Session I (top illumination),
- Domain B: Session II (bottom illumination).

These domains differed substantially in contrast, background appearance, and larval visual characteristics.

#### 3.7.2. CycleGAN Training

CycleGAN (Zhu et al., 2017) was trained using unpaired dish crops (256 × 256 px) sampled from both domains: 2,500 training and 500 test crops per domain, after automatic quality control based on brightness standard deviation. As an unpaired, unsupervised image-translation method, CycleGAN requires neither annotations nor paired correspondences between domains. The generator (ResNet with 9 residual blocks) and PatchGAN discriminator were trained for 200 epochs (λ_cycle = 10.0, λ_identity = 0.5) on a single NVIDIA H100 GPU.

The model optimized adversarial, cycle-consistency, and identity losses, enabling image translation while preserving larval morphology and spatial structure.

To avoid information leakage in the domain-adaptation evaluation, it is important to distinguish two distinct datasets. The unpaired crops used to train CycleGAN were unlabeled images used solely to learn the appearance mapping between illumination domains; CycleGAN never had access to the annotated detection labels. The evaluation of the adapted detector (Section 4.7) was performed strictly on the held-out, labeled Session II test split — the same frames used to quantify the cross-session baseline — with identical model weights, frames, labels, and evaluation parameters; the only difference between the baseline and adapted conditions was the CycleGAN style translation of the input images.

This test-time adaptation protocol ensures that the reported improvement reflects domain harmonization rather than additional supervision or label exposure.

#### 3.7.3. Domain Translation Experiments

Image translation was performed in both directions:

- A → B,
- B → A.

Translated images were used for model evaluation and domain adaptation experiments.

#### 3.7.4. Structural Fidelity Assessment

To verify preservation of biologically relevant information, transformed images were inspected visually and quantitatively. Larval shape integrity and centroid displacement were evaluated before and after domain translation.

### 3.8. Experimental Design

The following experimental configurations were evaluated:

1. Classical CV – Session I.
2. Classical CV – Session II.
3. YOLOv8 within-session evaluation.
4. YOLOv8 cross-session evaluation.
5. YOLOv8 with StyleGAN2-ADA augmentation.
6. YOLOv8 with CycleGAN-based domain adaptation.

This design enabled systematic assessment of tracking accuracy, domain-shift effects, and generalization performance.

### 3.9. Evaluation Metrics

Tracking Performance

Tracking quality was evaluated using:

- detection rate,
- missing frame rate,
- precision,
- recall,
- mAP@0.5.

Behavioral Agreement

Agreement between pipelines was assessed using:

- total distance traveled,
- average velocity,
- spatial coverage,
- heatmap similarity,
- Pearson correlation of group-level behavioral metrics.

Computational Performance

To support the comparative nature of this study, we also report computational characteristics of the pipelines, including model training environment, inference time per recording, and the effect of input resolution on throughput and detection completeness.

Deep learning models were trained on the WCSS computing cluster using NVIDIA H100 GPUs (batch size 8, imgsz = 640).

Cross-Session Generalization

Generalization performance was quantified as:

- performance drop between sessions,
- cross-session generalization was quantified as: performance drop between sessions; relative performance change after StyleGAN2-ADA augmentation; relative improvement after CycleGAN adaptation,
- relative improvement after CycleGAN adaptation.

### 3.10. Statistical Analysis

Descriptive statistics are reported as mean ± standard deviation unless stated otherwise. Agreement between behavioral outputs generated by the classical CV and YOLO pipelines was assessed at the level of group-mean traveled distance (n = 10 group-session means; five experimental groups × two sessions) using Pearson and Spearman correlation coefficients, the median pipeline ratio (YOLO/CV), and the mean absolute percentage error (MAPE). To quantify the uncertainty of these estimates, 95% confidence intervals (CIs) were obtained by non-parametric bootstrap resampling of the paired observations (10,000 iterations, percentile method). The systematic difference between pipelines was tested using the two-sided paired Wilcoxon signed-rank test and the paired t-test.

For detection-rate metrics, 95% bootstrap CIs were computed across the per-larva detection rates (n = 30 larvae per session, 10,000 iterations). Cross-session domain shift and the effect of CycleGAN-based adaptation were quantified as absolute and relative changes in detection and segmentation metrics evaluated on a fixed, identical set of test frames, ensuring direct comparability between conditions.

Statistical analyses were performed in Python (Python Software Foundation, 2021) using the SciPy and StatsModels libraries, with NumPy (Harris et al., 2020), pandas (McKinney, 2010), and Matplotlib (Hunter, 2007) employed for numerical processing and visualization.

## 4. Results

### 4.1. Performance of the Classical Computer Vision Pipeline

The classical computer vision (CV) pipeline achieved high tracking completeness across both recording sessions, confirming its suitability for controlled larval tracking experiments. The mean detection rate was 99.05% in Session I and 99.71% in Session II, corresponding to missing frame rates of 0.95% and 0.29%, respectively (Table 1). These results indicate that, after session-specific parameter tuning, the CV pipeline provided stable and nearly complete centroid trajectories.

**Table 1.** Baseline tracking performance of the classical CV pipeline.

| Metric | Session I | Session II |
| --- | --- | --- |
| Detection rate (%) | 99.05 | 99.71 |
| Missing frame rate (%) | 0.95 | 0.29 |

**Figure 1.**
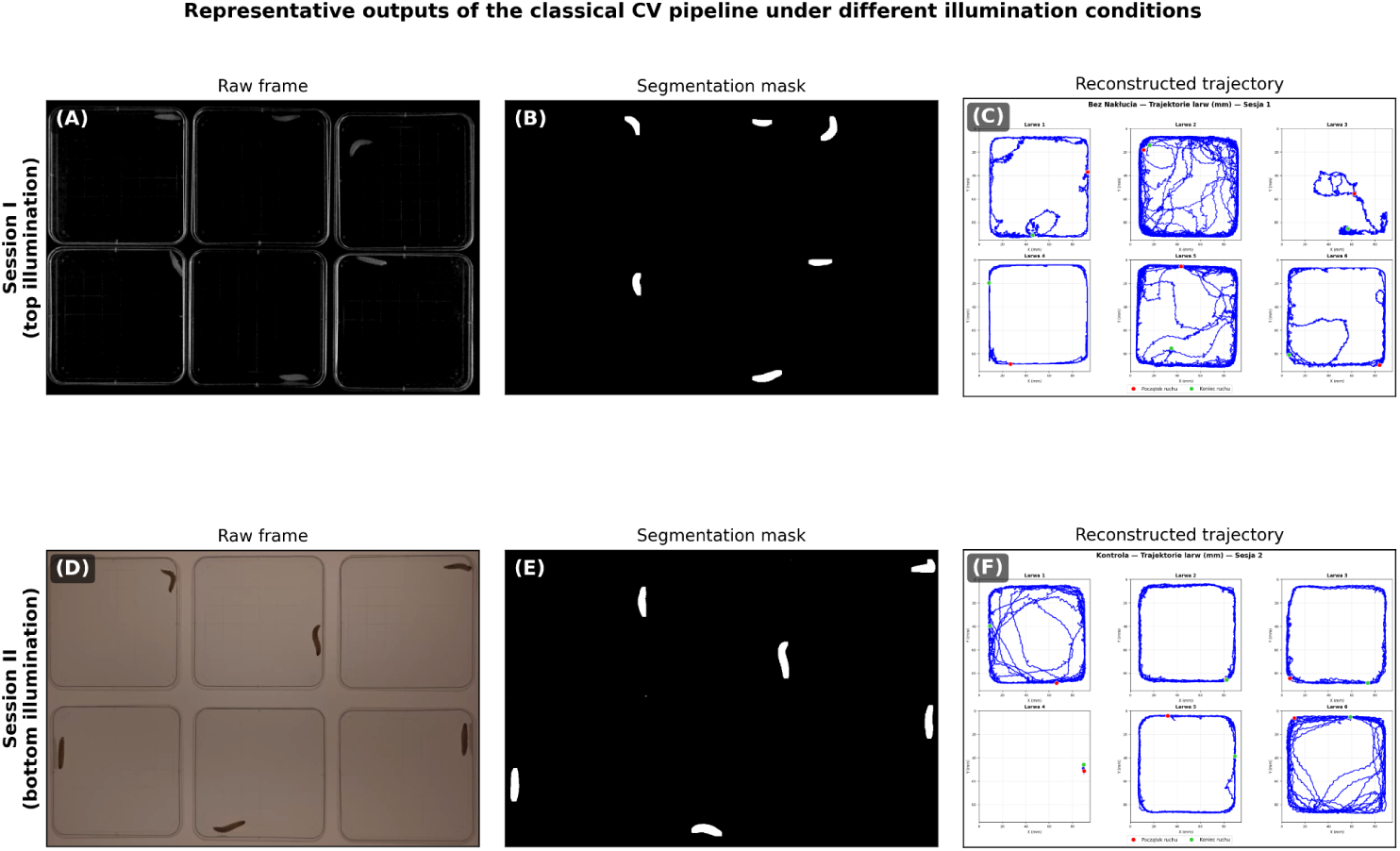
Representative outputs of the classical CV pipeline under different illumination conditions. (A–C) Session I (top illumination): raw frame, segmentation mask, and reconstructed trajectory. (D–F) Session II (bottom illumination): corresponding outputs.

Although the final detection rates were high in both sessions, this performance required session-specific preprocessing and manual adjustment of detection parameters. Thus, the CV pipeline served as a strong but manually tuned baseline.

### 4.2. Cross-Session Variability and Domain Shift

The two recording sessions differed substantially in illumination, background intensity, and larval appearance. These differences affected segmentation quality and required different preprocessing strategies. In the classical CV pipeline, this domain shift was reflected by the need for separate thresholding logic, contrast enhancement, and mask-parameter tuning between sessions.

Representative failure cases before and after preprocessing adjustment are shown in Fig. 2. Without appropriate correction, illumination artifacts produced fragmented contours or false detections. The introduction of CLAHE and the area-penalty contour selection mechanism improved segmentation stability and reduced artifact-driven tracking errors.

**Figure 2.**
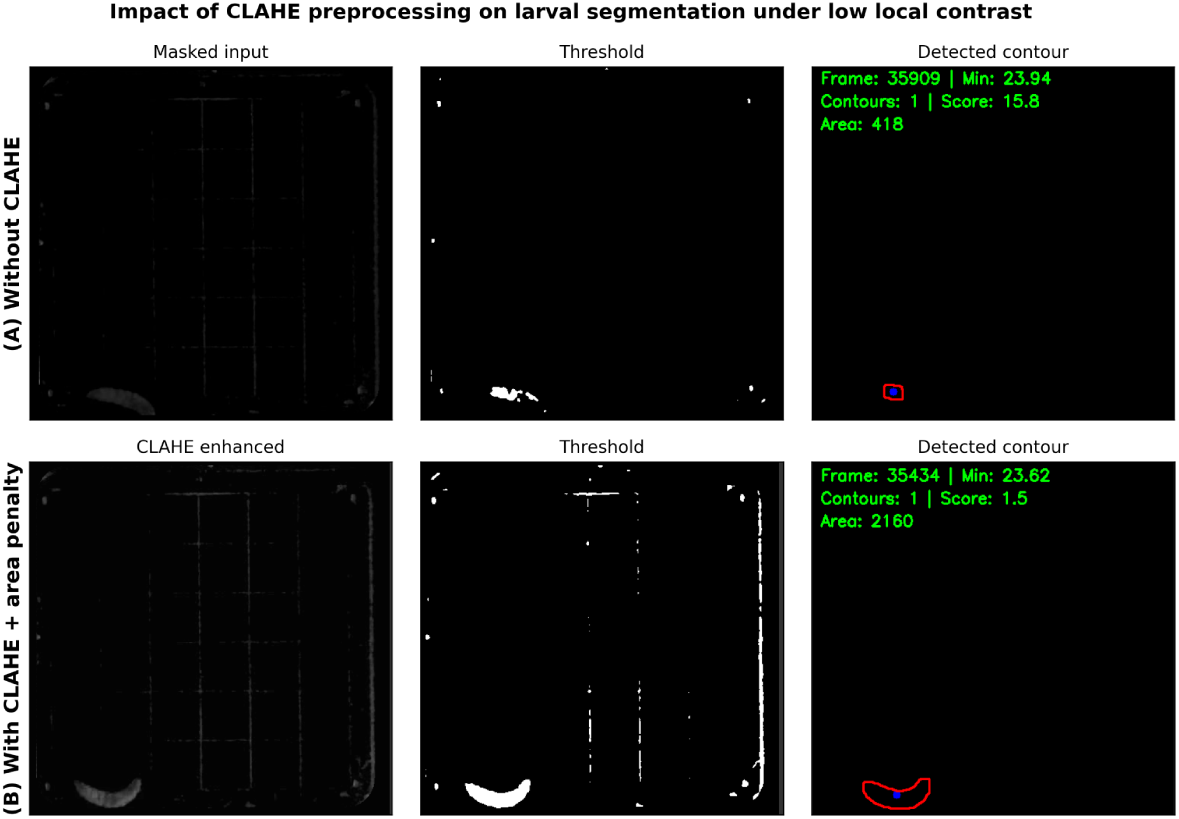
Effect of CLAHE preprocessing on segmentation in Session II. (A) Without CLAHE; (B) with CLAHE and area-penalty contour selection. Columns: masked input, threshold, detected contour.

These observations demonstrate that classical CV can be highly accurate in small-data biological settings, but its robustness depends on expert-driven adaptation to acquisition conditions.

### 4.3. Deep Learning Detection Performance

YOLOv8s-seg models achieved high within-session detection and segmentation performance. Three training configurations were evaluated: a combined model trained on both sessions, a Session I-only model, and a Session II-only model.

For bounding-box detection, mAP@0.5 ranged from 0.948 to 0.966 across models. For instance segmentation masks, mAP@0.5 ranged from 0.924 to 0.945 (Table 2)

**Table 2.**
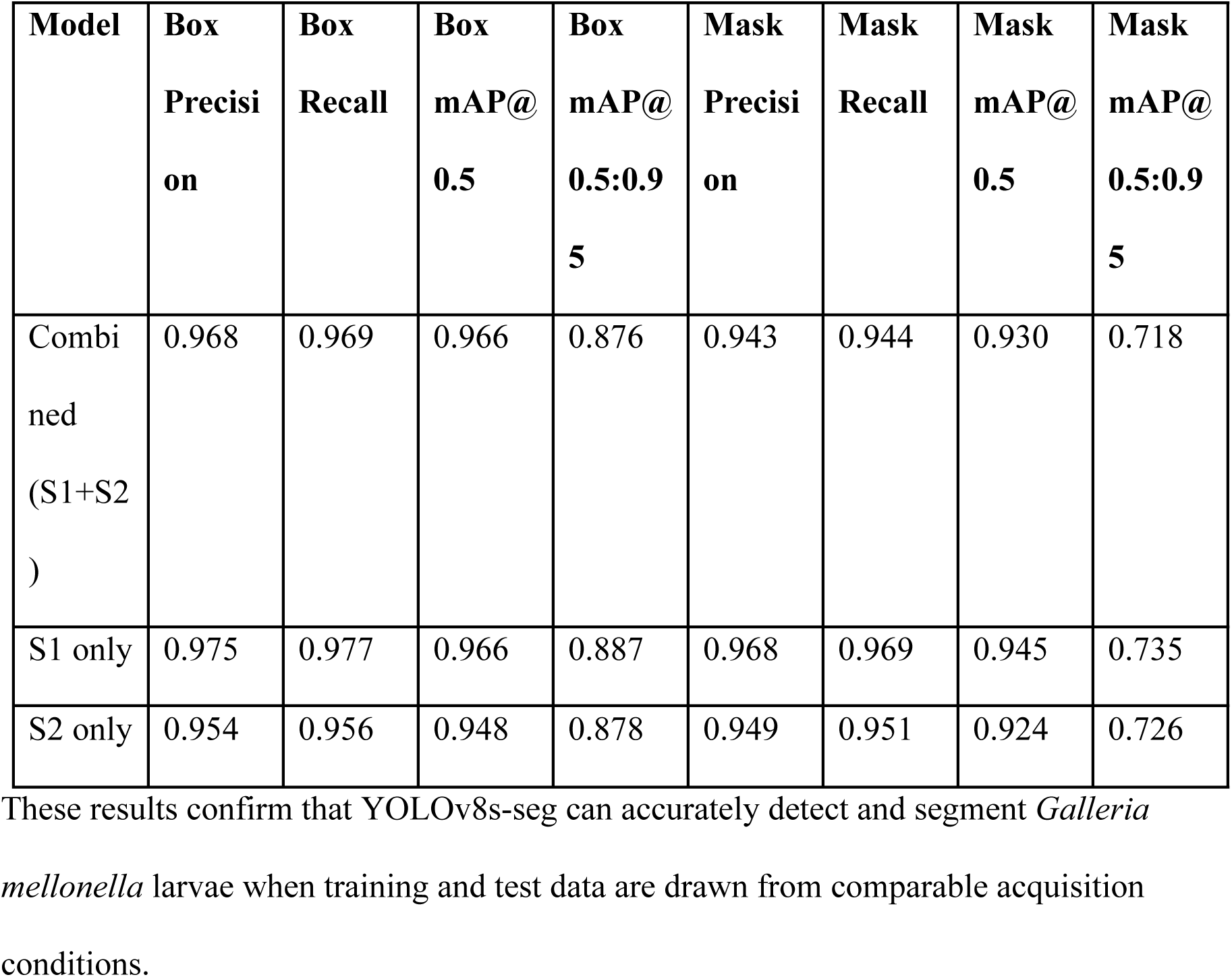
Within-session YOLOv8s-seg performance.

| Model | Box<br>Precision | Box<br>Recall | Box<br>mAP@<br>0.5 | Box<br>mAP@<br>0.5:0.95 | Mask<br>Precision | Mask<br>Recall | Mask<br>mAP@<br>0.5 | Mask<br>mAP@<br>0.5:0.95 |
| --- | --- | --- | --- | --- | --- | --- | --- | --- |
| Combined<br>(S1+S2) | 0.968 | 0.969 | 0.966 | 0.876 | 0.943 | 0.944 | 0.930 | 0.718 |
| S1 only | 0.975 | 0.977 | 0.966 | 0.887 | 0.968 | 0.969 | 0.945 | 0.735 |
| S2 only | 0.954 | 0.956 | 0.948 | 0.878 | 0.949 | 0.951 | 0.924 | 0.726 |

### 4.4. Cross-Session Generalization of YOLOv8s-seg

Despite strong within-session performance, cross-session evaluation revealed severe domain shift. When the model trained on Session I was tested on Session II, larval mask mAP@0.5 dropped to 0.091. The reverse transfer was even more severe: the model trained on Session II and tested on Session I achieved larval mask mAP@0.5 of only 0.00007 (Table 3).

**Table 3.** Cross-session performance for the larva class.

| Scenario | Train | Test | Box mAP@0.5 | Mask mAP@0.5 |
| --- | --- | --- | --- | --- |
| Within-session<br>S1 | S1 | S1 | 0.966 | 0.945 |
| Within-session<br>S2 | S2 | S2 | 0.948 | 0.924 |
| Cross-session<br>S1→S2 | S1 | S2 | 0.664 | 0.091 |
| Cross-session<br>S2→S1 | S2 | S1 | 0.049 | 0.00007 |
| Combined<br>baseline | S1+S2 | S1+S2 | 0.939 | 0.900 |

**Figure 3.**
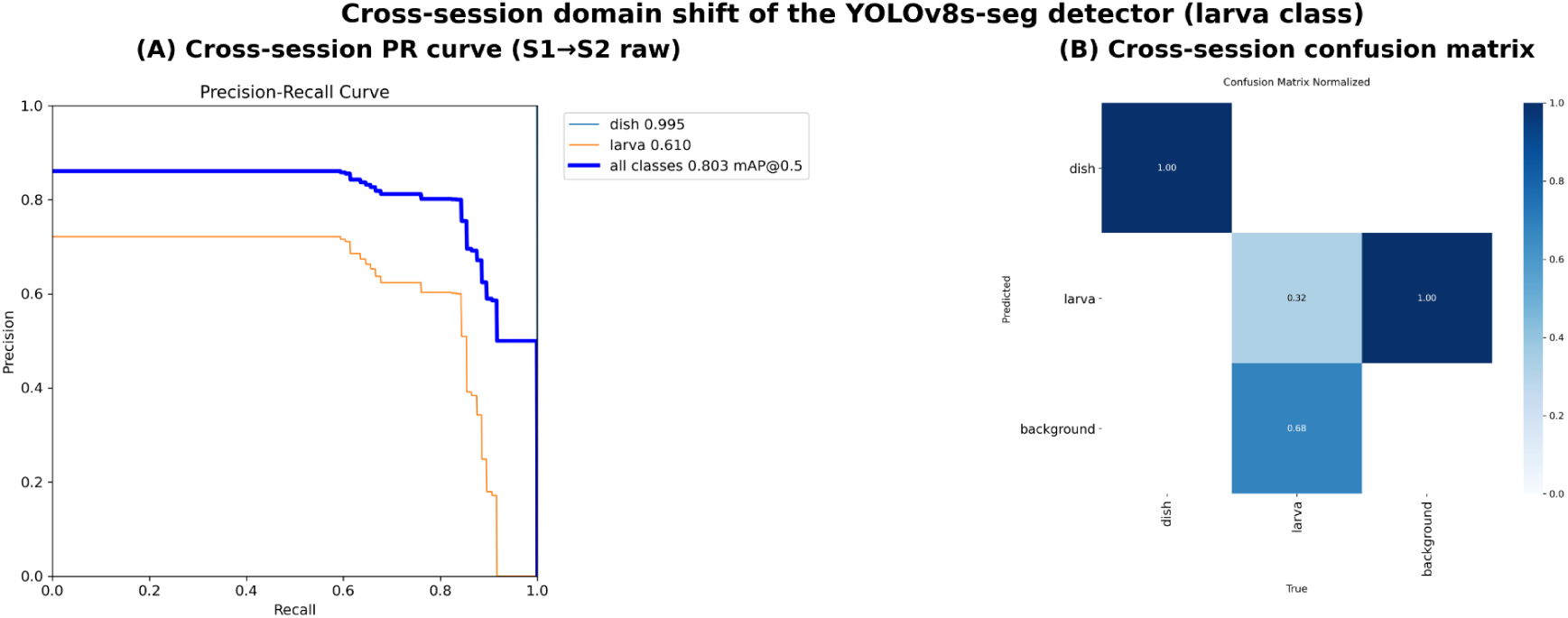
Cross-session domain shift of the YOLOv8s-seg detector (larva class). (A) Precision–recall curves (S1→S2). (B) Normalized confusion matrix.

### 4.5. Agreement Between OpenCV and YOLOv8s-seg + ByteTrack

The YOLOv8s-seg + ByteTrack pipeline enabled automated full-frame processing of six dishes simultaneously, eliminating manual dish isolation and per-dish parameter tuning. At the level of group-mean distance estimates, YOLO-derived trajectories showed strong agreement with the classical CV pipeline. The Pearson correlation between OpenCV and YOLO distance estimates was r = 0.89 (95% bootstrap CI [0.57, 1.00]; p = 0.0007), and the Spearman correlation was ρ = 0.94 (p < 0.0001). The median YOLO/OpenCV distance ratio was 0.99 (95% bootstrap CI [0.86, 1.05]) and the MAPE was 14.7%. Critically, the paired Wilcoxon signed-rank test revealed no statistically significant systematic difference between the two pipelines (W = 25.0, p = 0.85; paired t-test p = 0.95), indicating that the two methods produce statistically interchangeable group-level distance estimates rather than a consistent bias in either direction.

This agreement indicates that, when detection remains reliable, the deep learning pipeline reproduces group-level behavioral measurements obtained with the manually tuned classical CV approach. Thus, YOLOv8s-seg + ByteTrack provides a scalable alternative to OpenCV-based tracking, particularly when sufficient domain coverage is available during training. The relatively wide confidence interval for the Pearson coefficient reflects the modest number of group-level observations (n = 10) and should be interpreted accordingly.

### 4.6. Effect of StyleGAN2-ADA Generative Augmentation

StyleGAN2-ADA generated realistic synthetic images of Petri dishes and larvae, reproducing both dark-background and bright-background acquisition styles. The generator achieved an FID of approximately 32, indicating visually plausible image synthesis.

**Figure 4.**
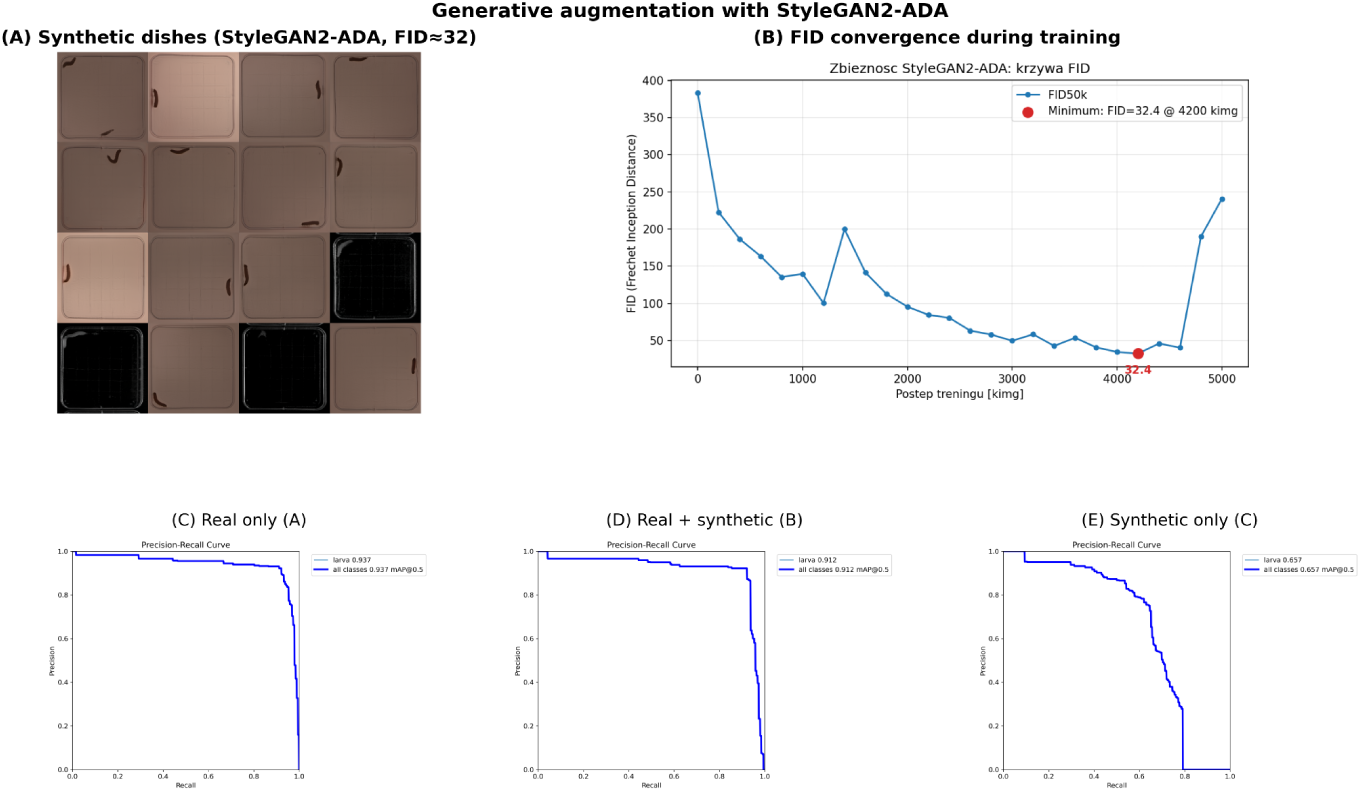
Generative augmentation with StyleGAN2-ADA. (A) Synthetic dishes (FID ≈ 32). (B) FID convergence. (C–E) Precision–recall curves: real only, real + synthetic, synthetic only.

Automatic pseudo-labeling of 1,500 generated images retained 886 high-confidence synthetic samples. These images were incorporated into the training set to evaluate whether generative augmentation improved larval detection.

Three configurations were compared: real-only training, real plus synthetic augmentation, and synthetic-only training (Table 4).

**Table 4.** Effect of StyleGAN2-ADA augmentation on larval detection.

| **Variant** | **Detection**<br>**rate (%)** | **Box**<br>**mAP@0.5** | **Box**<br>**mAP@0.5:**<br>**0.95** | **Mask**<br>**mAP@0.5** | **Mask**<br>**mAP@0.5:**<br>**0.95** |
| --- | --- | --- | --- | --- | --- |
| A —<br>Baseline,<br>real only | 86.1 | 0.937 | 0.807 | 0.834 | 0.508 |
| B — Real +<br>synthetic | 85.4 | 0.912 | 0.750 | 0.847 | 0.531 |
| C —<br>Synthetic<br>only | 17.2 | 0.657 | 0.383 | 0.664 | 0.277 |

Adding synthetic images to the real training set did not improve detection rate on full videos. The difference between the baseline and augmented models was minimal, with detection rates of 86.1% and 85.4%, respectively. Mask mAP@0.5 increased slightly from 0.834 to 0.847, but this did not translate into improved continuous tracking performance. The synthetic-only model achieved moderate mAP on static test frames but failed on full recordings, with a detection rate of only 17.2%. This result suggests that StyleGAN2-ADA captured relevant visual appearance features but did not sufficiently reproduce the full temporal and pose variability encountered during continuous video tracking.

It is important to note that the detection rates reported above (86.1% and 85.4%) reflect full-video tracking at an input resolution of imgsz = 640 px. The apparent discrepancy between high static-frame detection metrics (box mAP@0.5 in the 0.93–0.96 range) and the lower full-video detection rate is largely attributable to the small apparent size of the larva on the complete frame. After letterbox padding of the 16:9 recording to a square detection space, an individual larva occupies only approximately 15 px, which lies close to the lower size limit reliably resolved by the detector. Increasing the input resolution to imgsz = 1280 px enlarges the effective object size in the detection space and raises the full-video detection rate from approximately 85% to approximately 99%, without retraining or additional annotation. This indicates that the moderate detection rate observed at imgsz = 640 px is a resolution-related limitation rather than an intrinsic shortcoming of the detector, and that the generative-augmentation comparison above was performed under the more demanding low-resolution setting.

### 4.7. Effect of CycleGAN-Based Domain Adaptation

CycleGAN-based domain adaptation produced the strongest improvement in cross-session generalization. The most clinically and methodologically relevant experiment involved translating Session II test images into the visual style of Session I before applying the Session I-trained detector.

Before adaptation, the S1-trained YOLO model achieved only 0.078 mask mAP@0.5 on raw Session II images. After CycleGAN-based S2→S1 translation, mask mAP@0.5 increased to 0.884. Box mAP@0.5 increased from 0.610 to 0.884 (Table 5).

**Table 5.**
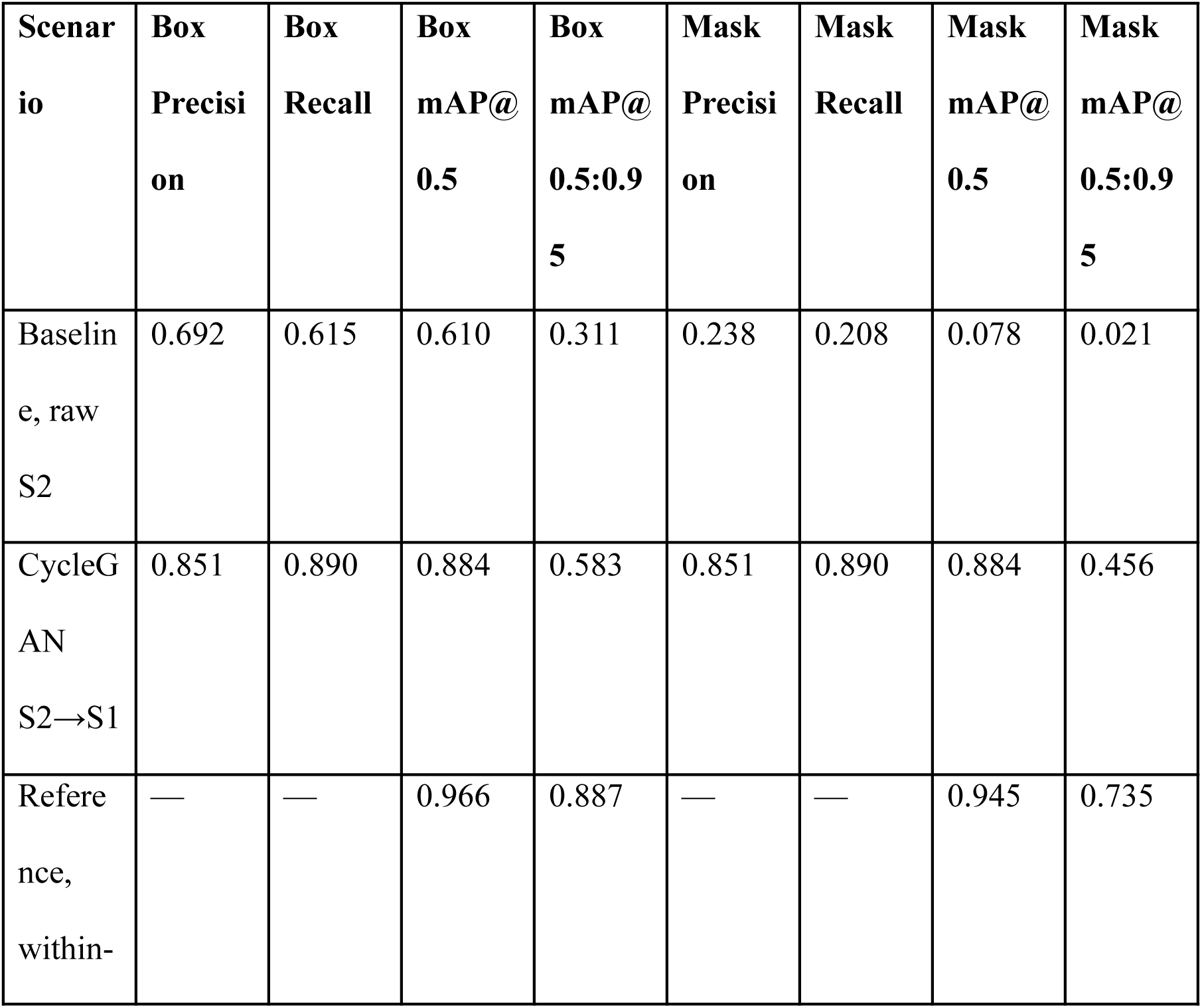
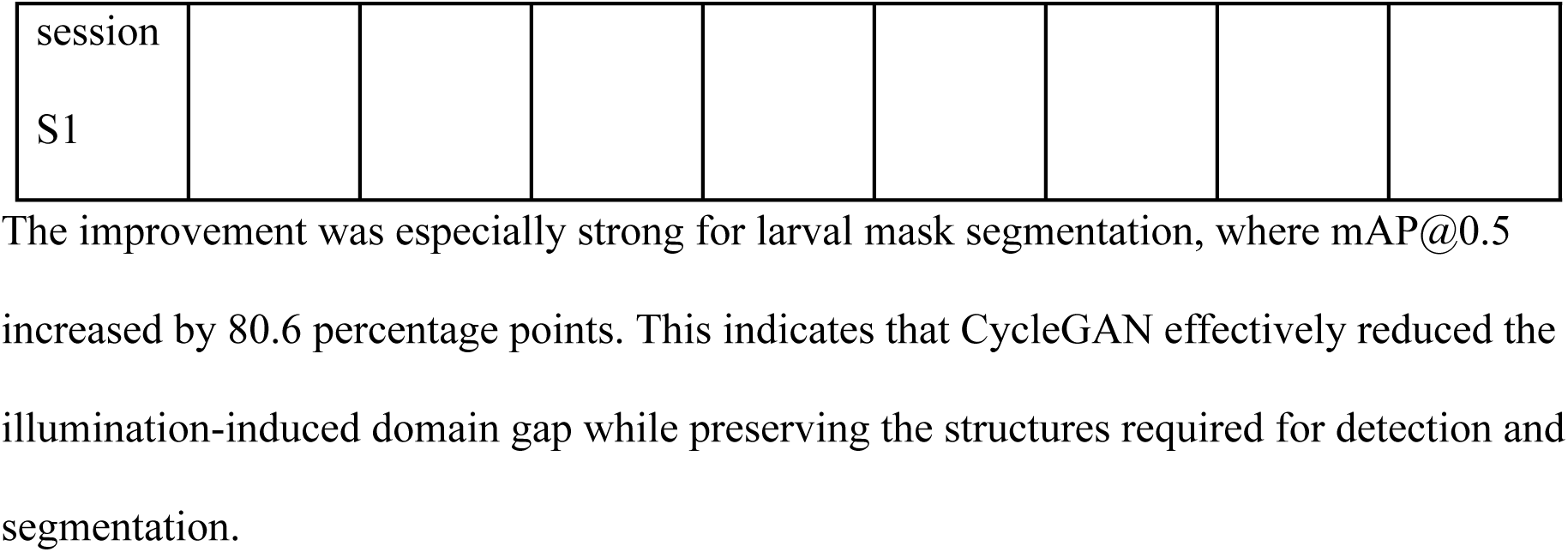
Effect of CycleGAN-based domain adaptation on cross-session larval detection.

The baseline raw-S2 values reported here (Box mAP@0.5 = 0.610; mask mAP@0.5 = 0.078) are slightly lower than the corresponding cross-session S1→S2 values in Table 3 (0.664 and 0.091). This difference arises because the present experiment was restricted to a fixed subset of 16 Session II test frames (split = test), identical to the set used for the adapted variant to ensure direct comparability, whereas the values in Table 3 were computed over a broader Session II evaluation set. The discrepancy therefore reflects the narrower test split rather than a methodological inconsistency, and both pairs of values confirm the same conclusion of near-complete failure of the S1-trained detector on raw Session II images.

**Figure 5.**
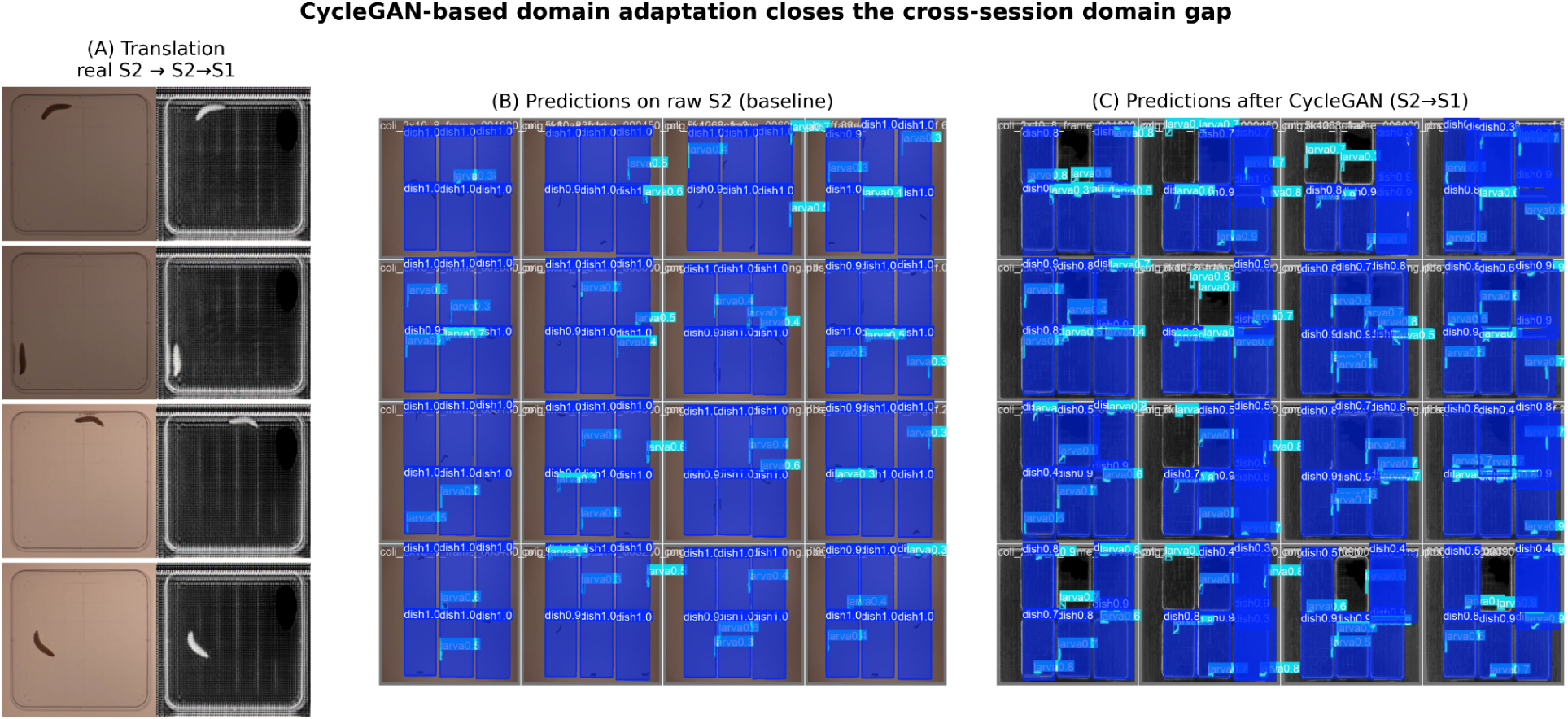
CycleGAN-based domain adaptation (S2→S1). (A) Sample translations (raw S2 vs. translated). (B) Detections on raw S2 (baseline). (C) Detections after CycleGAN translation.

Structural fidelity analysis further supported the validity of the adaptation. The median larval centroid drift after S2→S1 translation was 2.12 px, and 73.3% of image pairs showed centroid drift below 3 px. This suggests that the transformation primarily modified image appearance rather than larval geometry.

In contrast to the StyleGAN2-ADA experiments, where FID (≈32) provided a meaningful indicator of synthetic-image quality, FID was not adopted as the primary evaluation criterion for CycleGAN. Because image-to-image translation deliberately alters the global appearance of the source image (illumination, background intensity, and contrast), a relatively high FID between translated and target images (≈202) is expected and does not reflect a failure of the adaptation. For domain adaptation, structural-fidelity metrics that directly quantify preservation of biologically relevant information—most importantly the centroid drift of 2.12 px reported above—are more informative than distributional similarity scores, and were therefore used to assess CycleGAN translation quality.

### 4.8. Impact on Behavioral Representations

Detection quality directly influenced downstream behavioral analysis. Inaccurate or unstable detections affected trajectory reconstruction, traveled distance, velocity estimation, spatial occupancy maps, and directional movement patterns.

The classical CV pipeline provided nearly complete trajectories after manual tuning. The YOLOv8s-seg + ByteTrack pipeline produced comparable group-level distance estimates when detection was reliable, as shown by the strong OpenCV–YOLO agreement. However, under cross-session domain shift, degraded segmentation led to missing detections and unstable behavioral outputs.

The classical CV pipeline yielded biologically interpretable behavioral representations in both imaging sessions. For the majority of animals, reconstructed trajectories and occupancy heatmaps of the control group revealed consistent spatial patterns across sessions, including pronounced edge-associated movement along the Petri dish boundary. Two animals were excluded from this behavioral interpretation: larva 3 in Session I and larva 4 in Session II exhibited markedly anomalous patterns-erratic exploration and near-complete immobility (0.1% spatial coverage), respectively. These cases resulted from excessive needle puncture during handling rather than from intrinsic locomotor behavior, and are therefore treated as experimental noise. Across the remaining animals, the qualitative agreement between Session I (top illumination) and Session II (bottom illumination) confirms that, once reliable detection is achieved, the resulting behavioral descriptors are robust to the imaging conditions. CycleGAN adaptation contributes to this objective indirectly, by improving detection continuity under cross-session domain shift and thereby enabling stable behavioral reconstruction comparable to the manually tuned baseline.

**Figure 6.**
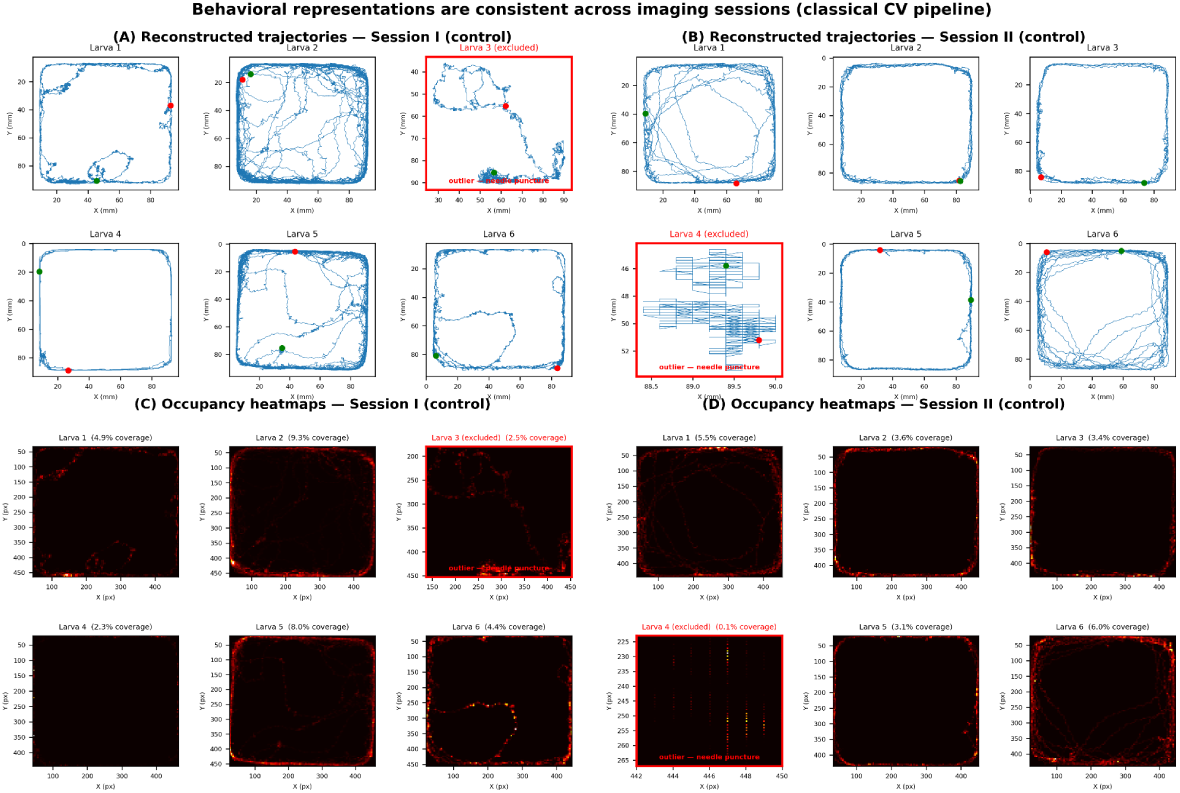
Behavioral representations obtained with the classical CV pipeline are consistent across imaging sessions. Reconstructed trajectories (A, B) and occupancy heatmaps (C, D) of the control group in Session I and Session II display consistent edge-associated movement along the Petri dish boundary. Larva 3 (Session I) and larva 4 (Session II) are technical outliers caused by excessive needle puncture (erratic exploration and near-complete immobility, respectively) and do not represent normal locomotor behavior. These findings demonstrate that improving cross-session detection performance is not only a technical objective but also a prerequisite for reliable quantitative behavioral phenotyping.

### 4.9. Qualitative Failure Analysis

Qualitative inspection supported the quantitative findings. Classical CV failure cases were mainly associated with illumination artifacts, reflections near dish borders, and fragmented contours. These errors were reduced by CLAHE preprocessing and the area-penalty contour selection mechanism.

YOLOv8s-seg performed well under within-session conditions but failed when larval appearance differed substantially from the training domain. In particular, segmentation masks were more vulnerable to domain shift than bounding boxes.

StyleGAN2-ADA generated visually realistic samples, but occasional artifacts were observed, including unrealistic larval duplication or insufficient representation of motion-related variability. CycleGAN improved visual consistency between sessions and substantially increased detection performance, although occasional minor structural distortions remained.

Overall, the results show that classical CV provides a strong interpretable baseline, YOLOv8 enables scalable full-frame automation, and CycleGAN-based adaptation effectively mitigates cross-session domain shift without additional manual annotation.

### 4.10. Computational Performance

Because computational cost is relevant for a comparative methodological study, we report the principal runtime characteristics of the two tracking pipelines (Table 6). Deep learning models were trained on the WCSS computing cluster using NVIDIA H100 GPUs; full YOLOv8s-seg training converged within early stopping (16–49 best epochs, depending on the scenario), and CycleGAN training required approximately 4.5 hours on a single H100. At inference, the classical CV pipeline required substantial human preparation time (manual ROI definition and per-recording parameter tuning, on the order of ∼30 minutes per recording) in addition to ∼5 minutes of processing per isolated dish. The YOLOv8s-seg + ByteTrack pipeline required essentially no manual preparation: a single script processed the full six-dish frame end-to-end. Single-threaded CPU inference took approximately 30 minutes per recording at imgsz = 640; on a CUDA-enabled GPU this is reduced by an estimated factor of 5–10× (∼3–6 minutes per recording).

**Table 6.** Computational characteristics of the tracking pipelines.

| Characteristic | Classical CV | YOLOv8s-seg +<br>ByteTrack |
| --- | --- | --- |
| Input | Isolated dish (1 larva) | Full frame (6 dishes) |
| Manual preparation | ~30 min/recording | None (automatic) |
| Per-recording parameters | Yes (10+ parameters) | No (single conf. threshold) |
| Detection rate | ~98–100% | ~85% (640 px) / ~99%<br>(1280 px) |
| Inference time | ~5 min/dish | ~30 min/recording (CPU)* |
| Training hardware | — | NVIDIA H100 (WCSS<br>cluster) |

*\*Measured in single-threaded CPU mode; GPU (CUDA) inference is estimated to be 5–10× faster (~3–6 min/recording).*

Input resolution governs a clear accuracy–throughput trade-off. At imgsz = 640 an individual larva spans only ∼15 px in the letterboxed detection space, yielding a mean full-video detection rate of 84.9% (95% bootstrap CI [83.1, 86.7]; per-session: S1 85.3% [82.6, 87.8], S2 84.4% [81.9, 86.9]). Doubling the input resolution to imgsz = 1280 enlarged the effective larval size to ∼30 px and raised the mean detection rate to ∼99% (S1 99.5%, S2 98.9%) without retraining, at the cost of increased computation and greater bounding-box jitter. Group-level kinematic results were therefore reported at imgsz = 640, which provided the most stable agreement with the classical pipeline.

## 5. Discussion

The present study investigated the robustness of automated behavioral tracking in *Galleria mellonella* larvae under heterogeneous imaging conditions and evaluated whether generative artificial intelligence can improve cross-session generalization. Three principal findings emerged from our experiments. First, a carefully designed classical computer vision pipeline achieved remarkably high tracking completeness despite the relatively simple methodology. Second, deep learning models demonstrated excellent within-session performance but experienced severe degradation when exposed to previously unseen acquisition conditions. Third, CycleGAN-based domain adaptation substantially reduced the domain gap and restored detection performance without requiring additional manual annotation.

### 5.1. Classical Computer Vision Remains Highly Competitive in Small-Data Biological Applications

One of the most notable findings of this study is the strong performance of the classical computer vision pipeline. Detection rates exceeded 99% in both recording sessions, demonstrating that carefully engineered image-processing workflows remain highly effective in controlled biological environments. These findings are particularly relevant because contemporary behavioral analysis increasingly relies on deep learning solutions, often under the assumption that neural network-based approaches universally outperform traditional computer vision techniques.

Our results suggest that this assumption does not necessarily hold in small-data experimental settings. The larval tracking task investigated here involved a relatively constrained environment consisting of a single animal per Petri dish, fixed camera geometry, and limited background complexity. Under such conditions, classical methods benefited from transparency, interpretability, and low computational requirements while achieving performance comparable to more complex learning-based approaches.

At the same time, this strong performance must not be misinterpreted as evidence that classical computer vision is universally superior to deep learning. The high accuracy of the classical pipeline was achieved only at the cost of substantial expert effort and manual engineering. Reliable detection required session-specific tuning of threshold values, contrast-enhancement procedures (CLAHE), morphological-operation parameters, and contour-selection criteria, and in our workflow the masking and threshold parameters were adjusted individually for each of the larva-recordings. This manual tuning is precisely what does not generalize: the pipeline is not a turnkey solution but a carefully hand-crafted procedure whose parameters are specific to a given acquisition setup and whose successful operation presupposes operator expertise. Each new illumination configuration, dish type, or camera geometry would in general require renewed manual calibration.

Three limitations therefore characterize the classical approach and should be stated explicitly: (i) it requires manual, per-recording parameter tuning; (ii) it is not a general-purpose detector and does not transfer automatically to new acquisition conditions; and (iii) it depends on domain expertise to be configured and validated. In contrast, the deep learning pipeline, once trained, processed full-frame recordings of six dishes with a single confidence threshold and no per-recording tuning, and — when domain coverage is adequate or restored through domain adaptation — produced behavioral measurements statistically interchangeable with the manually tuned baseline. The appropriate conclusion is thus not that one paradigm dominates the other, but that classical CV offers an interpretable, low-data baseline at the price of manual labor and limited scalability, whereas deep learning offers automation and scalability at the price of annotated data and sensitivity to domain shift, the latter being addressable by generative domain adaptation.

### 5.2. Domain Shift Represents the Primary Limitation of Deep Learning-Based Behavioral Tracking

While YOLOv8s-seg achieved excellent within-session detection and segmentation accuracy, cross-session experiments revealed a dramatic decline in performance. The severe reduction in segmentation accuracy observed when models were evaluated on unseen illumination conditions demonstrates that domain shift remains a critical challenge for automated behavioral phenotyping.

Importantly, this degradation occurred despite the relatively simple nature of the experimental environment. The two recording sessions differed primarily in illumination configuration and image contrast, yet these differences were sufficient to reduce larval segmentation performance by more than an order of magnitude. This observation is consistent with broader findings in computer vision and biomedical imaging, where deep learning models frequently learn domain-specific visual characteristics in addition to task-relevant features.

The results emphasize that high within-dataset performance should not be interpreted as evidence of robust generalization. In biological research, where data are often collected across multiple experimental sessions, instruments, and laboratories, models may encounter imaging conditions that differ substantially from those represented during training. Consequently, evaluation protocols should include cross-session validation whenever possible to avoid overly optimistic estimates of model performance.

### 5.3. Generative Augmentation Provides Limited Benefits for Cross-Session Robustness

The use of StyleGAN2-ADA for synthetic image generation produced visually realistic larval images (FID ≈ 32) and enabled the creation of additional training data. However, the impact of generative augmentation on downstream detection performance was negligible: adding 886 high-confidence synthetic images to the real training set changed the full-video detection rate only marginally (86.1% → 85.4%), a difference far smaller than the between-individual variability of the detection rate (62.7%–98.9%). We interpret this negative result as informative rather than as a failure of generative augmentation per se, and several converging explanations help to clarify why synthetic data did not improve robustness.

First, and most importantly, generative augmentation primarily addresses data scarcity, whereas the limiting factor in our setting was domain shift, not training-set size. The baseline detector already operated near the performance ceiling on within-domain static frames (box mAP@0.5 in the 0.93–0.96 range), leaving little headroom for additional in-domain data to exploit. Synthetic images sampled from the same learned distribution as the real training data add appearance variability within the existing domains but do not introduce the out-of-domain illumination characteristics responsible for the cross-session gap. Augmentation that does not span the target domain cannot close a domain gap, regardless of its visual quality.

Second, the contrast between the synthetic-only model’s moderate static-frame performance (box mAP@0.5 = 0.657) and its collapse on full recordings (detection rate = 17.2%) reveals a deeper limitation: visual realism of individual frames does not guarantee coverage of the full operational distribution. Continuous tracking confronts the detector with tens of thousands of frames spanning motion blur, transitional poses, partial self-occlusion, and edge-of-dish configurations. StyleGAN2-ADA, trained on temporally subsampled static crops, reproduces the marginal appearance of typical larvae convincingly but under-represents these rarer dynamic and pose-related states. A low FID — a distributional similarity measure dominated by common, easy samples — is therefore a poor predictor of usefulness for a tracking task whose failures concentrate in the tails of the distribution.

Third, residual generator artifacts (e.g., occasional duplication of larvae within a single dish, inconsistent with the one-larva-per-dish protocol) introduce label noise during pseudo-labeling that can partially offset any benefit of the additional samples.

Taken together, these observations indicate that generative augmentation is most useful when training data are genuinely the bottleneck, and that it is unlikely to resolve domain-shift problems on its own. This also explains why CycleGAN, which explicitly targets the appearance gap between domains rather than expanding in-domain diversity, succeeded where StyleGAN2-ADA did not (Section 5.4). A practical methodological corollary is that detectors intended for continuous tracking should be evaluated on full recordings, since static-frame mAP can substantially overestimate their real-world utility.

### 5.4. CycleGAN Effectively Mitigates Cross-Session Domain Shift

Among all evaluated approaches, CycleGAN-based domain adaptation produced the most substantial improvement in cross-session performance. Translating images from the target acquisition domain into the visual style of the source domain dramatically increased detection and segmentation accuracy while preserving larval morphology and trajectory structure.

This finding supports the hypothesis that the primary source of performance degradation was appearance-related rather than structural. By harmonizing illumination characteristics and image contrast between sessions, CycleGAN effectively transformed the testing distribution into a representation more closely aligned with the training data distribution. Importantly, structural fidelity analysis demonstrated that image translation introduced only minimal centroid displacement, indicating preservation of biologically relevant spatial information. This observation is particularly important for behavioral phenotyping applications, where even small geometric distortions could potentially influence downstream movement metrics.

The substantial improvement achieved without additional manual annotation highlights one of the major advantages of unsupervised domain adaptation. In many biological studies, collecting new annotations for every acquisition condition is impractical or prohibitively time-consuming. CycleGAN-based adaptation therefore represents a promising strategy for improving reproducibility and scalability of behavioral tracking workflows.

### 5.5. Consequences for Behavioral Phenotyping

An important contribution of this study is the demonstration that tracking accuracy directly influences downstream behavioral interpretation. Errors in segmentation and centroid localization propagated to higher-level behavioral descriptors, including traveled distance, velocity estimates, and spatial occupancy patterns.

The strong agreement observed between OpenCV-derived and YOLO-derived behavioral metrics under favorable conditions suggests that multiple tracking methodologies can produce biologically comparable results when detection performance is adequate. However, this agreement deteriorates when domain shift causes unstable detection, highlighting the importance of evaluating behavioral pipelines beyond conventional computer vision metrics.

The preservation of trajectory structures and occupancy heatmaps following CycleGAN adaptation further suggests that improvements in detection performance translate into more reliable behavioral measurements. Consequently, domain adaptation should be viewed not merely as a technical optimization procedure but as an essential step toward robust quantitative phenotyping.

A further practical consideration concerns input resolution. The moderate full-video detection rate observed for the YOLOv8s-seg + ByteTrack pipeline was strongly influenced by the small apparent size of the larva on the full frame: at imgsz = 640 px an individual larva spans only about 15 px after letterbox padding. Raising the input resolution to imgsz = 1280 px recovered detection rates close to those of the classical CV pipeline (from approximately 85% to approximately 99%), demonstrating that this limitation is readily addressable at inference time. This distinction is methodologically important, because it shows that low full-video detection rates should not be conflated with poor model quality (static-frame mAP remained high); rather, they reflect the trade-off between processing resolution and computational cost when tracking small organisms on wide-field recordings.

### 5.6. Limitations and Future Directions

Several limitations should be acknowledged. First, the study was conducted using a single species and a relatively controlled experimental setup. Additional studies involving different organisms, recording systems, and behavioral paradigms will be required to establish the generalizability of the proposed framework.

Second, the deep learning experiments were based on a relatively small manually annotated dataset. Although this reflects realistic constraints encountered in many biological laboratories, larger datasets may improve both detection performance and robustness.

Third, only a single domain adaptation architecture was evaluated. Recent developments in diffusion models, adversarial adaptation frameworks, and foundation vision models may provide additional opportunities for improving cross-domain generalization.

Future work should therefore investigate multi-laboratory datasets, larger annotation resources, and alternative adaptation strategies, while also exploring the integration of behavioral classification and anomaly-detection models built upon the proposed tracking framework.

### 5.7. Conclusions

This study demonstrates that cross-session domain shift represents a major obstacle to reliable automated behavioral tracking in biological experiments. While classical computer vision achieved excellent performance after manual tuning and deep learning models provided highly accurate within-session detection, both approaches exhibited limitations when confronted with heterogeneous acquisition conditions. CycleGAN-based domain adaptation emerged as the most effective solution, substantially improving cross-session generalization while preserving biologically meaningful behavioral information. These findings highlight the importance of explicitly addressing domain shift in behavioral phenotyping pipelines and support the use of generative domain adaptation as a practical strategy for enhancing robustness and reproducibility in small-data biological imaging applications.

## Data Availability Statement

The datasets generated and analyzed during the current study consist of video recordings of *Galleria mellonella* larvae acquired under two experimental illumination conditions, together with manually annotated image subsets used for deep learning model training and evaluation. The datasets are available from the corresponding author upon reasonable request.

## Ethics Statement

*Galleria mellonella* larvae are invertebrate organisms and are not subject to animal experimentation regulations applicable to vertebrate animals in many jurisdictions, including the European Union Directive 2010/63/EU. All experimental procedures were performed in accordance with institutional guidelines for handling invertebrate model organisms.

No additional experiments were conducted specifically for the purpose of this computational study. The analyses reported herein were performed on video recordings acquired as part of previously completed laboratory experiments.

## Code Availability Statement

The source code used for video preprocessing, classical computer vision-based tracking, trajectory reconstruction, behavioral feature extraction, YOLOv8s-seg training and evaluation, ByteTrack-based trajectory association, StyleGAN2-ADA augmentation experiments, CycleGAN domain adaptation, and statistical analyses is publicly available at: https://github.com/kasjansmigielski/cv-dl-larvae-domain-adaptation

The repository contains scripts necessary to reproduce the principal analyses presented in this study, including data preprocessing, model training, evaluation, and behavioral metric extraction.

## Acknowledgements

The biological studies using the *Galleria mellonella* model, including larval culture, experimental design, larval injection, video recording, and raw data acquisition, were conducted at the Division of Translational Technologies, Wroclaw Medical University. The resulting preliminary data were provided to the authors of this publication as a scientific courtesy for further analysis and preparation of the manuscript.

## Funding

The larval husbandry and preparation, behavioral recordings, preliminary data acquisition was supported by the Wroclaw Medical University grant no. SUBZ.D300.26.018.

## Declaration of Generative AI Use

During the preparation of this manuscript, the main and corresponding author used Claude Opus 4.6 (Anthropic) for the purpose of language polishing and grammatical correction of the text. The output was thoroughly reviewed, verified, and edited by the corresponding author (K.Ś.) to ensure accuracy and to confirm that no misrepresentations of the original content occurred. The authors take full responsibility for the content of the published work.

## Conflict of Interest

The authors declare that they have no known competing financial interests or personal relationships that could have appeared to influence the work reported in this paper.

## CRediT Author Contributions

Kasjan Śmigielski Conceptualization; Methodology; Software; Validation; Formal analysis; Investigation; Data curation; Writing – Original Draft; Visualization

Natalia Piórkowska Visualization; Writing – Original Draft; Writing – Review & Editing; Project Administration

## Equations

Equation (1). Pixel-to-millimeter conversion: pos_mm = pos_px · s, s = 0.20 mm/px

Equation (2). Total traveled distance: D = Σ (t=1 → T−1) √[(x□₊₁ − x□)² + (y□₊₁ − y□)²] · s

Equation (3). Average velocity: v̅ = D / T_total = D / (T / FPS)

Equation (4). Spatial coverage: C = (N_visited / N_total) · 100%

Equation (5). Movement direction angle (compass convention): θ□ = (atan2(−Δx, Δy) · 180/π) mod 360, where Δx = x□₊₁ − x□ and Δy = y□₊₁ − y□

## Notes

https://github.com/kasjansmigielski/cv-dl-larvae-domain-adaptation

### Competing Interest Statement

The authors have declared no competing interest.

